# Recommendations for sample pooling on the Cepheid GeneXpert^®^ system using the Cepheid Xpert^®^ Xpress SARS-CoV-2 assay

**DOI:** 10.1101/2020.05.14.097287

**Authors:** Michael G. Becker, Tracy Taylor, Sandra Kiazyk, Dana R. Cabiles, Adrienne F.A. Meyers, Paul A. Sandstrom

## Abstract

The coronavirus disease 2019 (Covid-19) pandemic, caused by SARS-CoV-2, has resulted in a global testing supply shortage. In response, pooled testing has emerged as a promising strategy that can immediately increase testing capacity. Here, we provide support for the adoption of sample pooling with the point-of-care Cepheid Xpert^®^ Xpress SARS-CoV-2 molecular assay. Corroborating previous findings, the Xpert^®^ Xpress SARS-CoV-2 assay limit of detection was comparable to central laboratory reverse-transcription quantitative PCR tests with observed SARS-CoV-2 detection below 100 copies/mL. The Xpert^®^ Xpress assay detected SARS-CoV-2 after samples with minimum viral loads of 461 copies/mL were diluted into six sample pools. Based on these data, we recommend the adoption of pooled testing with the Xpert^®^ Xpress SARS-CoV-2 assay where warranted by population public health needs. The suggested number of samples per pool, or pooling depth, is unique for each point-of-care test site and should be determined by assessing positive test rates. To statistically determine appropriate pooling depth, we have calculated the pooling efficiency for numerous combinations of pool sizes and test rates. This information is included as a supplemental dataset that we encourage public health authorities to use as a guide to make recommendations that will maximize testing capacity and resource conservation.

## 1. Introduction

The coronavirus disease 2019 (COVID-19) pandemic has caused an unprecedented demand for global testing supplies. In response, public health officials are searching for innovative ways to increase testing capacity in the face of limited resources. One approach that could be rapidly deployed to achieve increased SARS-CoV-2 testing capacity is pooled sample testing. The number of samples to be combined into each test pool, or pooling depth, is determined by test sensitivity and community disease prevalence, with some laboratories pooling up to 10 (1), 30 (2), or 48 (3) samples using the Corman quantitative reverse transcription PCR (RT-qPCR) test (4). Similar strategies should be explored for currently deployed SARS-CoV-2 point-of-care tests, such as the Cepheid *<u>Xpert</u>*<u>^®^</u> *<u>Xpress SARS-CoV-2 assay</u>*.

The Cepheid *<u>Xpert</u>*<u>^®^</u> *<u>Xpress SARS-CoV-2 assay</u>* is a rapid, fully-automated, and self-contained multiplex qualitative RT-qPCR test for SARS-CoV-2 detection. The Cepheid *<u>Xpert Xpress SARS-CoV-2 assay</u>* targets two regions of the SARS-CoV-2 genome: the N (nucleocapsid) region and the E (envelope) region. The test is interpreted as positive for SARS-CoV-2 if either of the two analytes produce a Ct below 45. The test is performed on the Cepheid GeneXpert system in single-use cartridges, with an approximate run time of 50 minutes. This Cepheid *<u>Xpert</u>*<u>^®^</u> *<u>Xpress SARS-CoV-2 assay</u>* received approval from Health Canada on March 24, 2020 under interim order authorization. Evaluation of the Cepheid SARS-CoV-2 assay is ongoing, with higher reported sensitivity than the Abbott *<u>ID</u> <u>Now SARS-CoV-2 Assay</u>* (5, 6), and high agreement (>99%) with the Roche Cobas 6800 system (5, 7, 8) and the Centers for Disease Control and Prevention (CDC) RT-qPCR test (8). Using viral recombinants to contrive samples, Cepheid reports 100% sensitivity (n=35) at 250 copies (cp)/mL. Using synthetic RNA controls, Zhen et al. (6) reported 100% sensitivity at 100 cp/mL (n=10) and a 87.5% sensitivity at 50 cp/mL (n=8).

Given the high sensitivity of the *<u>Xpert</u>*<u>^®^</u> *<u>Xpress SARS-CoV-2 assay</u>*, it is reasonable to propose that it could be suitable for pooled testing. Here, the potential for pooled SARS-CoV-2 testing was assessed on the GeneXpert system using a small panel of clinical specimens with low-to mid-range viral loads that were diluted with known clinical negative samples. The results here corroborate previous findings that the LOD for the Cepheid test is likely <100 cp/mL. Additionally, data generated by this study suggest that the GeneXpert device can be effectively applied for SARS-CoV-2 pooled sample testing in pools containing up to at least six individual samples. Finally, a reference dataset is provided that can be used by public health authorities to advise point-of-care test sites on the optimal number of samples to combine per pool given their current positive test rates.

## 2. Materials and Methods

### 2.1 Viral Culture

High-titre inactive SARS-CoV-2 culture (Strain VIDO; GISAID Accession: EPI_ISL_425177), made inactive by gamma-irradiation, was provided by the Special Pathogens Program of the National Microbiology Laboratory. Briefly, virus was cultured in Vero cells in minimum essential media and cellular debris were removed via low-speed centrifugation. Nominal viral load of the inactivated virus was determined by quantitation of dilutions that fell within another standard curve that was developed using serial dilutions of the *<u>SeraCare AccuPlex</u>*<u>™</u> *<u>SARS-CoV-2 Reference Material Kit</u>* (0505-0126). The *<u>AccuPlex</u>*<u>™</u> *<u>SARS-CoV-2 Reference Material Kit</u>* consists of recombinant Sindbis virus containing SARS-CoV-2 RNA, at a concentration of 5000 cp/mL.

### 2.2 Clinical Specimens

Clinical nasopharyngeal swab samples were collected in 1 mL of viral transport media, and provided by Cadham Provincial Laboratory (CPL; Winnipeg, Canada). All provided samples were previously tested and characterized using their approved SARS-CoV-2 diagnostic RT-qPCR assay. The ethics-exempt panel used for this study consisted of six positive CPL clinical samples and an additional two low viral load swab samples (Ct=37/Ct=38) provided by the Influenza and Respiratory Viruses Program of the National Microbiology Laboratory. Pooled negative samples were also provided by the Influenza and Respiratory Viruses Program.

### 2.3 Standard Curve for the Xpert Xpress^®^ SARS-CoV-2 assay

A standard curve was produced to facilitate the quantitation of SARS-CoV-2 viral load for each of the clinical samples used in this study. To produce the curve, 10-fold serial dilutions of inactivated high-titre SARS-CoV-2 were prepared in viral transport media to yield a series from 6 × 10^8^ cp/mL to 6 × 10^0^ cp/mL. For each step of the dilution series, 300 µL was pipetted into an <u>Xpert^®^ Xpress SARS-CoV-2</u> cartridge. The linear equation from the standard curves for analytes E and N were used to determine the nominal viral load of undiluted clinical samples following testing with the *<u>Xpert^®^ Xpress SARS-CoV-2 assay</u>*. The reported value is the average between the N and E targets.

### 2.4 Sample Pool Tests

All testing was performed with the same GeneXpert system used to produce the standard curve, and all replicates of a given sample were performed on the same system module. Each sample was tested without pooling on the GeneXpert system and Ct values were used to determine nominal viral load using the standard curve. Each sample was then diluted in a pool of confirmed negative clinical specimens to simulate either three-sample pools (three-fold dilution) or six-sample pools (six-fold dilution). To conserve the <u>Xpert^®^ Xpress SARS-CoV-2 assay</u> cartridges for clinical use, only six-sample pools were performed in triplicate and a limited number of samples were included in our panel. Each pool was created individually to account for errors in pipetting.

### 2.5 Calculation of Pooling Efficiency

To provide guidance on pooling efficiency, the impact of pooling on testing capacity was calculated at various pooling depths (1-10) and individual positive test rates (0-100%). Similar to the statistical approach used by Hanel and Thurner (9), a custom Python script was used to calculate the pooled testing capacity relative to non-pooled testing capacity, represented as a percentage, for each combination. A value above 100% indicates that testing capacity has increased, whereas values below 100% indicate decreased capacity. To calculate relative testing capacity, for each combination of pool sizes and positive test rates, the proportion of pools that would be SARS-CoV-2 positive (*P*^*S+*^) were determined with the following equation:

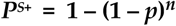

where *n* is the pool size and *p* is the proportion of individual tests that are positive. The average number of tests needed per pool was then determined by multiplying the proportion of pools that are positive by the pool size to determine the cost of retests, in addition to the original test that was needed for the pool itself. The average number of tests (*T*) needed to process each pool is therefore determined by:

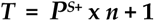

Finally, relative testing capacity was calculated by dividing the average number of tests required for each pool, divided by the number of samples tested:

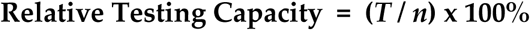

## 3. Results

### 3.1 The Xpert^®^ Xpress SARS-CoV-2 assay can be used to provide quantitative results

Although the <u>Xpert^®^ Xpress SARS-CoV-2 assay</u> is considered a qualitative test, it does provide output Ct values that can be used to approximate viral loads using a standard curve. To produce a standard curve, 10-fold serial dilutions of inactivated high-titre SARS-CoV-2 were prepared in viral transport media from 6 × 10^8^ cp/mL to 6 × 10^0^ cp/mL. All dilutions above 6 × 10^1^ were recorded as SARS-CoV-2 positive by the assay (Supplemental Table 1), consistent with the previously observed LOD (6) for the <u>Xpert^®^ Xpress SARS-CoV-2 assay</u> of <100 cp/mL. The resulting curve was highly linear (R^2^ > 0.999), suggesting that the Ct values can be used for quantitation across the entire range of our standard curve. The qPCR efficiency for the E and N analytes was 99.8% and 96.6%, respectfully (Figure 1).

**Table 1:** Five clinical samples collected at the Cadham Provincial Laboratory (CPL) were selected for analysis with Ct values ranging from 23-35 as determined by the CPL in-house RT-qPCR test. Each sample was tested with the *Xpert*^®^_*Xpress SARS-CoV-2 assay* as an undiluted sample, diluted three-fold in negative clinical samples to simulate a three sample pool, and diluted six-fold in negative clinical samples to simulate a six sample pool (performed in triplicate). Ct values are provided for the envelope (E), nucleocapsid (N), and sample processing control (SPC) targets at each dilution. Nominal viral load was determined from the standard curve using the results from the *Xpert*^®^_*Xpress SARS-CoV-2 assay* using undiluted clinical specimens. For the six sample pool replicates, standard deviation was calculated for each target.

| Sample ID | RT-qPCR Ct Value | Nominal Viral Load (cp/mL) | Undiluted |  |  | Three Sample Pool |  |  | Six Sample Pool (Replicate 1) |  |  | Six Sample Pool (Replicate 2) |  |  | Six Sample Pool (Replicate 3) |  |  | Replicate Standard Dev. |  |  |
| --- | --- | --- | --- | --- | --- | --- | --- | --- | --- | --- | --- | --- | --- | --- | --- | --- | --- | --- | --- | --- |
|  |  |  | E | N2 | SPC | E | N2 | SPC | E | N2 | SPC | E | N2 | SPC | E | N2 | SPC | E | N2 | SPC |
| CPL1 | 23 | 2,452,553 | 22.2 | 24.7 | 29.2 | 22.8 | 24.3 | 24.3 | 24.3 | 26.2 | 27.3 | 23.8 | 25.8 | 27.6 | 24.3 | 26.5 | 27.3 | 0.21 | 0.29 | 0.14 |
| CPL2 | 26 | 154,663 | 26.1 | 28.9 | 28.7 | 27.1 | 28.0 | 28.0 | 28.0 | 30.6 | 27.8 | 28.2 | 30.9 | 28.1 | 28.0 | 30.8 | 27.7 | 0.09 | 0.12 | 0.17 |
| CPL3 | 31 | 6439 | 30.5 | 33.9 | 28.4 | 31.9 | 33.5 | 33.5 | 33.5 | 37.4 | 28.1 | 34.1 | 36.7 | 27.9 | 33.5 | 36.6 | 28.2 | 0.37 | 0.36 | 0.12 |
| CPL4 | 33 | 2245 | 32.0 | 35.5 | 28.6 | 33.3 | 35.6 | 35.6 | 35.6 | 37.7 | 27.4 | 33.0 | 36.1 | 27.7 | 35.6 | 38.7 | 27.7 | 1.07 | 1.07 | 0.14 |
| CPL5 | 35 | 938 | 33.6 | 36.3 | 28.0 | 36.2 | 40.7 | 40.7 | 40.7 | 41.1 | 27.6 | 39.0 | 39.2 | 27.6 | 40.7 | 39.1 | 27.7 | 1.77 | 0.92 | 0.05 |

**Figure 1:**
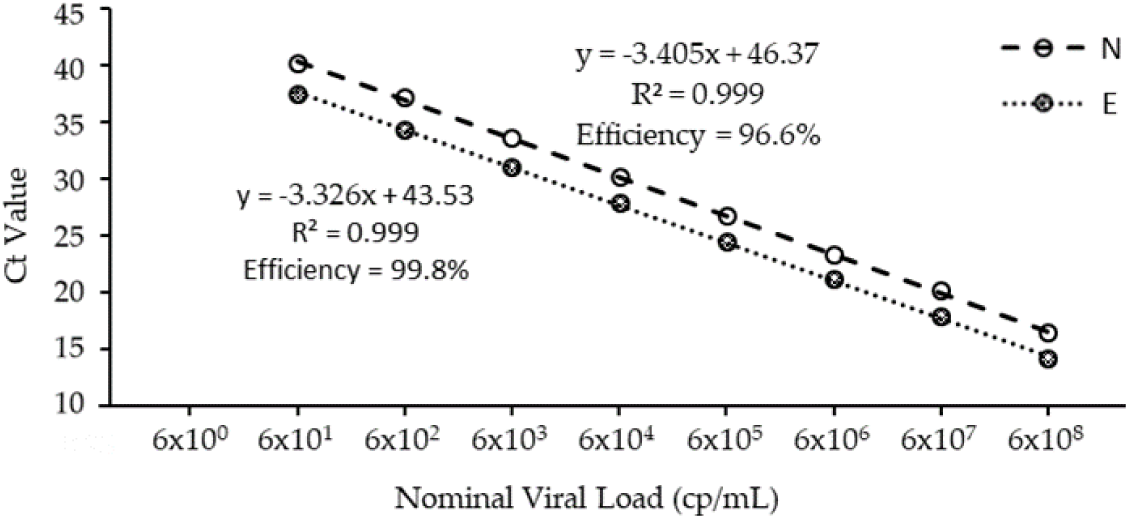
Standard curve for the *Xpert*^®^_*Xpress SARS-CoV-2 assay* nucleocapsid (N; empty circle with a dashed line) and envelope (E; filled circle with a dotted line) targets. Curve was produced with serially-diluted gamma-irradiated virus culture (GISAID Accession: EPI_ISL_425177) produced at the National Microbiology Laboratory.

### 3.2 Determining the effect of sample pooling on the Xpert^®^ Xpress SARS-CoV-2 assay

To determine the effect of sample pooling on the sensitivity of the <u>Xpert^®^ Xpress SARS-CoV-2 assay</u>, five clinical samples were selected with Ct values ranging from 23-35 as determined by the Corman RT-qPCR test performed at the Cadham Provincial Laboratory in Winnipeg, MB. Each sample was first individually tested without pooling on the GeneXpert^®^ system, and resultant Ct values were converted to nominal viral loads using the standard curve. Input viral loads ranged from approximately 938 cp/mL to 2.85 million cp/mL (Table 1). Each of these samples was then diluted in confirmed SARS-CoV-2 negative clinical samples to simulate three sample or six sample pooling. Although Ct values were higher, as anticipated after dilution in a pool, the Xpert^®^ <u>Xpress SARS-CoV-2 assay</u> correctly identified each pool qualitatively as SARS-CoV-2 positive. Standard deviation of Ct values between replicates increased at high Ct values, likely due to sampling and PCR biases. Though not clinically relevant, this would likely affect accurate quantitation of viral load at higher Ct values.

To better observe the effects of sample pooling near the <u>Xpert^®^ Xpress SARS-CoV-2 assay</u>’s LOD, an additional three clinical samples were selected with high Ct values (>37). This included a discordant sample that was not detected by the Cadham Provincial Lab RT-qPCR test, but was subsequently identified as weakly positive on the GeneXpert (CT=43.5/39.2). At initial viral loads of 461 and 1362 cp/mL, the <u>Xpert^®^ Xpress SARS-CoV-2 assay</u> detected SARS-CoV-2 after six-fold pooling with negative specimens, while our weak positive (64 cp/mL) returned a negative result (Table 2). Additionally, the E target was not detected in one of the pools; however, only one detected analyte is needed to return an actionable positive test result.

**Table 2:** An additional three clinical specimens with high Ct values were selected to better observe the effect of sample pooling close to the limit of detection of the *Xpert*^®^_*Xpress SARS-CoV-2 assay*. This included two samples provided by the National Microbiology Laboratory and one from the Cadham Provincial Laboratory (CPL), which is a discordant sample not detected by CPL’s Corman RT-qPCR test, but detected as a weak positive by the Xpert^®^ assay. At the six-fold dilution, the weak positive was no longer detected by the assay.

| Sample ID | RT-qPCR Ct Value | Nominal Viral Load (cp/mL) | Undiluted |  |  | Six Sample Pool |  |  |
| --- | --- | --- | --- | --- | --- | --- | --- | --- |
|  |  |  | E | N2 | SPC | E | N2 | SPC |
| NML1 | 37 | 1362 | 32.8 | 36.1 | 27.8 | 38.9 | 39.2 | 27.8 |
| NML2 | 38 | 461 | 34.9 | 37.1 | 29.4 | ND* | 38.9 | 27.7 |
| CPL6 | ND* | 64† | 43.5 | 39.2 | 28.3 | ND* | ND* | 28.2 |

\* ND; Not Detected
† Ct value was outside of the standard curve (E) and viral load was inferred through extrapolation

### 3.3 Determining the optimal pool size

An objective of this study is to provide guidance for when sample pooling is a viable option for SARS-CoV-2 testing with the *<u>Xpert^®^ Xpress SARS-CoV-2 assay</u>*, or any sensitive SARS-CoV-2 test in general. At high positive testing rates, pooling may actually increase the number of tests required to screen samples and increase turnaround time. Further, deciding what pooling depth to use is arbitrary without understanding the relationship between pooling depths and positive test rates. We determined testing capacity using various combinations of pool sizes (1-10) and test rates (0-100% in increments of 0.1%). A complete summary of all combinations can be found in Supplemental Table 2, and a graphical representation of a subset of these data is shown in Figure 2.

**Figure 2:**
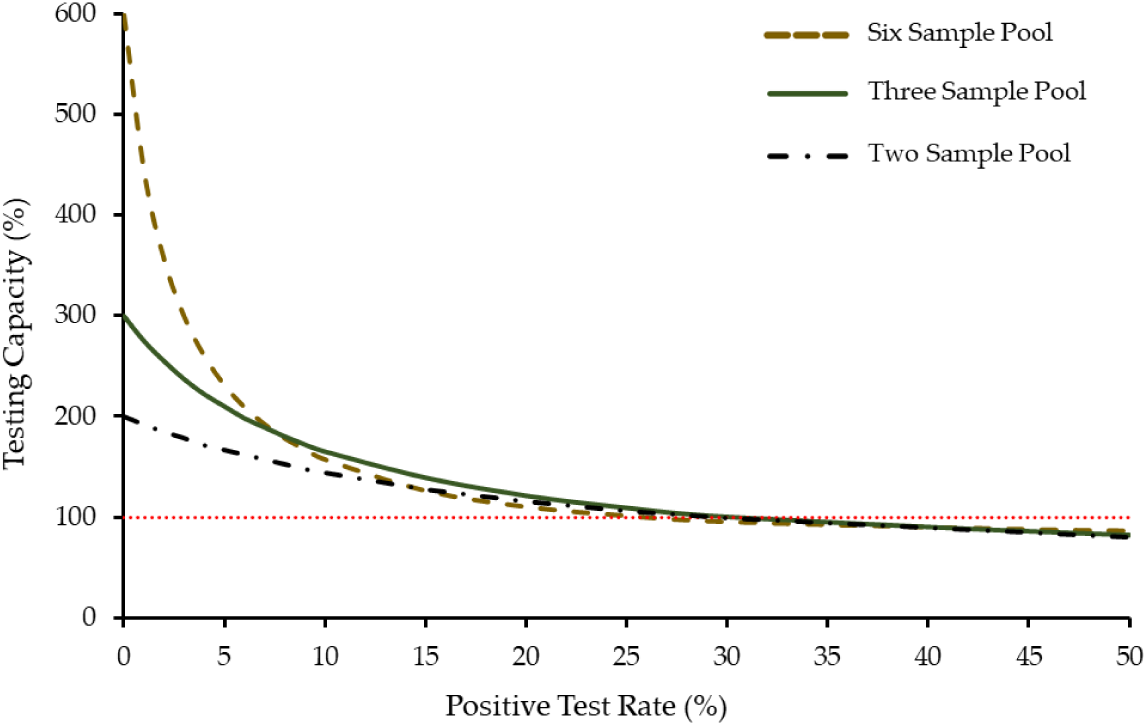
The effect of sample pooling on testing capacity using pools of two (black dashed-dotted line), three (green solid line), or six (brown dashed line) samples. For each pool size, testing capacity is plotted against the rate of positive individual tests. The red dotted line represents the point at which pooled testing decreases capacity and is no longer viable. The cross-over point, when three sample pooling is more efficient than six sample pooling, occurs at 7.6%.

This information can help medical authorities provide informed recommendations pertaining to sample pooling. For example, no pooling strategy is effective when positive test rates exceed ∼30%. Additionally, at no combination of pool size and positive test rates is two sample pooling more efficient than three sample pooling (Figure 2). Although at low positive test rates aggressive pooling is favored, this quickly changes when test rates increase above 1%. For example, if the positive test rate at a site is ∼3% the ideal pool size would be six samples.

## 4. Discussion

The results of this study strongly suggest that sample pooling is a viable option for SARS-CoV-2 testing using the *<u>Xpert^®^ Xpress SARS-CoV-2 assay</u>*. All samples tested positive after pooling, except for a high-Ct discordant positive with a nominal viral load of 64 cp/mL. At this level of sensitivity, pooled tests should detect SARS-CoV-2 in the vast majority of clinical cases; a study following 80 patients at different stages of infection detected average sample viral loads of >10^4^ from 1 day before to 7 days after disease onset, using sputum (n=67), throat (n=42), and nasal (n=1) swabs (10), with the lowest observed viral load of 641 copies/mL. Another research group determined average viral loads to be >10^5^ at the onset of mild to moderate symptoms (11). When testing asymptomatic individuals, results still show typical Ct values of 22-31 with the Corman RT-qPCR assay (12–14).

One challenge that may prevent some point-of-care testing sites from adopting a pooled testing strategy is the lack of mechanical pipettes. The *<u>Xpert^®^ Xpress SARS-CoV-2 assay</u>* is provided with single-use transfer pipettes that dispense 300 µL of sample. With small pool sizes, multiple samples can be combined into a 5 mL specimen tube or 15 mL canonical tube, and inverted to mix. Subsequently 300 µL of this pool can be transferred into a test cartridge. With this approach, pooled testing with the *<u>Xpert^®^ Xpress SARS-CoV-2 assay</u>* could be readily achieved in a resource-limited setting with the provision of additional 300 µL transfer pipettes.

Several other strategies for SARS-CoV-2 pooled testing are being investigated such as combinatorial testing, or matrix testing (3, 15). In this approach, samples are combined into multiple pools, such that each sample is tested multiple times across multiple pools. The combination of SARS-CoV-2 positive pools can identify individual positives with limited retesting required. Although this strategy is promising, it works best for high-throughput laboratories processing batches of hundreds of samples using 96- or 384-well plates and real-time PCR machines. Because of the need for larger batch sizes and its more complicated testing design, a combinatorial approach is unlikely to be feasible with point-of-care tests that perform only a few tests in a single run, such as the *<u>Xpert^®^ Xpress SARS-CoV-2 assay</u>*.

Another pooling strategy proposed by the German Red Cross Blood Donor Service and Geothe University is swab pooling, or the mini-pool method (16). Multiple swabs can be combined into a single tube at the point of collection, rather than the traditional method of pooling transport media or extracted RNA. As a result, there is minimal loss of sensitivity as no dilution is occurring. The main disadvantage of this approach is that the pooled samples need to be collected simultaneously and at the same location; however, the swab pool approach could be applied in certain scenarios. For example, this strategy may be appropriate for door-to-door household testing, workplace screening, or its intended of purpose of blood donor screening. This approach could easily be combined with traditional pooling to substantially increase testing capacity with the *<u>Xpert^®^ Xpress SARS-CoV-2 assay</u>* or other validated molecular test method.

## 5. Conclusions

This study provides a resource that can be used to determine the appropriate pool size to use at each testing site. Public health authorities can approximate positive tests rates, and use this information with the reference table (Supplemental Table 2) to make appropriate recommendations on pooling strategies. The application of sample pooling, when possible, can be used to immediately increase testing capacity on the GeneXpert^®^ system while conserving limited resources. Future experiments should investigate if more extensive pooling is viable on the GeneXpert^®^ system, similar to the aggressive pooling strategies being explored for the laboratory-based RT-qPCR tests.

## 6. Acknowledgements

The authors would like to acknowledge partners who provided the clinical specimens used in this study. This includes the Cadham Provincial Laboratory of Manitoba, the Special Pathogens Program of the National Microbiology Laboratory, and the Influenza and Respiratory Viruses Program of the National Microbiology Laboratory.

## 7. Contributions

M.G.B., P.S., and A.F.A.M. conceptualized the study. All authors contributed to study design and methodology. M.G.B. and T.T. prepared the manuscript. M.G.B. analyzed the data and prepared the figures. M.G.B., T.T., S.K., and D.C. performed the experiments. All authors edited and approved the manuscript for submission.

## 8. Conflicts of Interest

The authors declare no conflicts of interest.

## Notes

### Competing Interest Statement

The authors have declared no competing interest.

### Summary of Updates

This version of the manuscript has been revised to include supplemental datasets.

