## Supplemental Table 1 for "Recommendations for sample pooling on the Cepheid GeneXpert^®^ system using the Cepheid Xpert^®^ Xpress SARS-CoV-2 assay"

**Supplemental Table 1:** Serial dilutions of high-titre irradiated SARS-CoV-2 tested with the <u>*Xpert Xpress SARS-CoV-2 assay*</u>. Results show Ct values of the envelope (E), nucleocapsid (N), and sample processing control (SPC) targets at each dilution.

| Approx. viral concentration (cp/mL) | Result | E | N | SPC |
|---|---|---|---|---|
| $6 \times 10^0$ | Negative | ND* | ND* | 28.3 |
| $6 \times 10^1$ | Positive | 37.5 | 40.2 | 28.3 |
| $6 \times 10^2$ | Positive | 34.3 | 37.2 | 27.9 |
| $6 \times 10^3$ | Positive | 31.1 | 33.6 | 28.3 |
| $6 \times 10^4$ | Positive | 27.9 | 30.2 | 28 |
| $6 \times 10^5$ | Positive | 24.5 | 26.7 | 28.1 |
| $6 \times 10^6$ | Positive | 21.2 | 23.3 | 27.5 |
| $6 \times 10^7$ | Positive | 17.8 | 20.2 | 28.3 |
| $6 \times 10^8$ | Positive | 14.1 | 16.4 | 27.8 |

*ND = Not Detected
